# PLK1- and PLK4-mediated asymmetric mitotic centrosome size and positioning in the early zebrafish embryo

**DOI:** 10.1101/2020.04.13.039362

**Authors:** LI Rathbun, AA Aljiboury, X Bai, J Manikas, JD Amack, JN Bembenek, H Hehnly

## Abstract

Factors that regulate mitotic spindle positioning have been elucidated *in vitro*, however it remains unclear how a spindle is placed within the confines of extremely large cells. Our studies identified a uniquely large centrosome structure in the early zebrafish embryo (246.44±11.93μm^2^ mitotic centrosome in a 126.86±0.35μm diameter cell), whereas *C. elegans* centrosomes are notably smaller (6.75±0.28μm^2^ mitotic centrosome in a 55.83±1.04μm diameter cell). During early embryonic cell divisions, cell size changes rapidly in *C. elegans* and zebrafish embryos. Notably, mitotic centrosome area scales closely with changing cell size compared to changes in spindle length for both organisms. One interesting difference between the two is that mitotic centrosomes are asymmetric in size across embryonic zebrafish spindles, with the larger mitotic centrosome being 2.14±0.13-fold larger in size than the smaller. The largest mitotic centrosome is placed towards the embryo center in a Polo-Like Kinase (PLK) 1 and PLK4 dependent manner 87.14±4.16% of the time. We propose a model in which uniquely large centrosomes direct spindle placement within the disproportionately large zebrafish embryo cells to orchestrate cell divisions during early embryogenesis.

## RESULTS AND DISCUSSION

During early embryogenesis, rapid cell divisions increase the number of cells in an embryo to ensure that proper tissue and organ formation can proceed during later development. However, it remains unclear how the mitotic spindle is able to position itself within the confines of a cell that is disproportionately large. Previous studies have proposed that spindle size scales to cell size in embryos [1,2]. However, this poses the question whether the mitotic spindle adapts to the rapidly changing cell size during early embryonic cell divisions. This study aims to understand the previously unknown mechanism by which cell division is regulated during early development in extremely large cells.

One proposed model is that large embryonic cells use acentrosomal microtubule nucleation sites so that astral microtubules can reach the cortex in large cells [3]. The mitotic centrosome/spindle pole assembles the microtubule-based spindle, and one spindle pole consists of two centrioles surrounded by pericentriolar material (PCM) that contains microtubule nucleation sites [4]. Typically, astral microtubules emanate from the centrosome and project towards the cell cortex, where they anchor and facilitate pulling forces to position the spindle and undergo cell division [4]. In the proposed acentrosomal model, microtubule nucleation sites exist outside of the centrosome through branched microtubules, positioning astral microtubules closer to the cell cortex in large cells [3]. However, we find that large dividing zebrafish embryo cells have notably large mitotic centrosomes that scale with cell size. We hypothesize that large centrosomes are used to assist astral microtubules in reaching the cortex in large cells.

We present a model where mitotic centrosome size scales with cell size, and that this scaling requires Polo-Like Kinase (PLK) 1 and PLK4.

### Mitotic centrosome area scales with cell length during embryonic cell divisions in *C. elegans* and zebrafish

We used embryos from the invertebrate *Caenorhabditis elegans* (*C. elegans)* and vertebrate *Danio rerio* (zebrafish) as model systems to identify whether mitotic centrosomes scale with cell size. These organisms were chosen based on their stark differences in size, embryo morphology, and organism complexity (Supplementary Figure 1A-D). *C. elegans* embryos develop within an eggshell, where early divisions occur asynchronously [5] (Figure 1A, Supplementary Figure 1B-C, Supplementary Video 1). In contrast, early zebrafish embryos undergo rapid cleavage stage cell divisions on top of a yolk (Figure 1B, Supplementary Figure 1B, D, Supplementary Video 1). The first ten cell divisions occur synchronously, before transitioning to an asynchronous wave [6]. During the first five cell divisions, blastomeres create a cellular monolayer on top of the yolk. Each division during this stage occurs perpendicular to the plane of the previous division, leading to the construction of a monolayer grid (model, Figure 1B) [6]. This is clearly visualized through the use of a fluorescent microtubule transgenic zebrafish line (EMTB-3xGFP) [7], where the 16-cell stage embryo contains mitotic spindles oriented perpendicular to the previous division at the 8-cell stage (Figure 1B, Supplementary Video 2-3).

**Figure 1.**
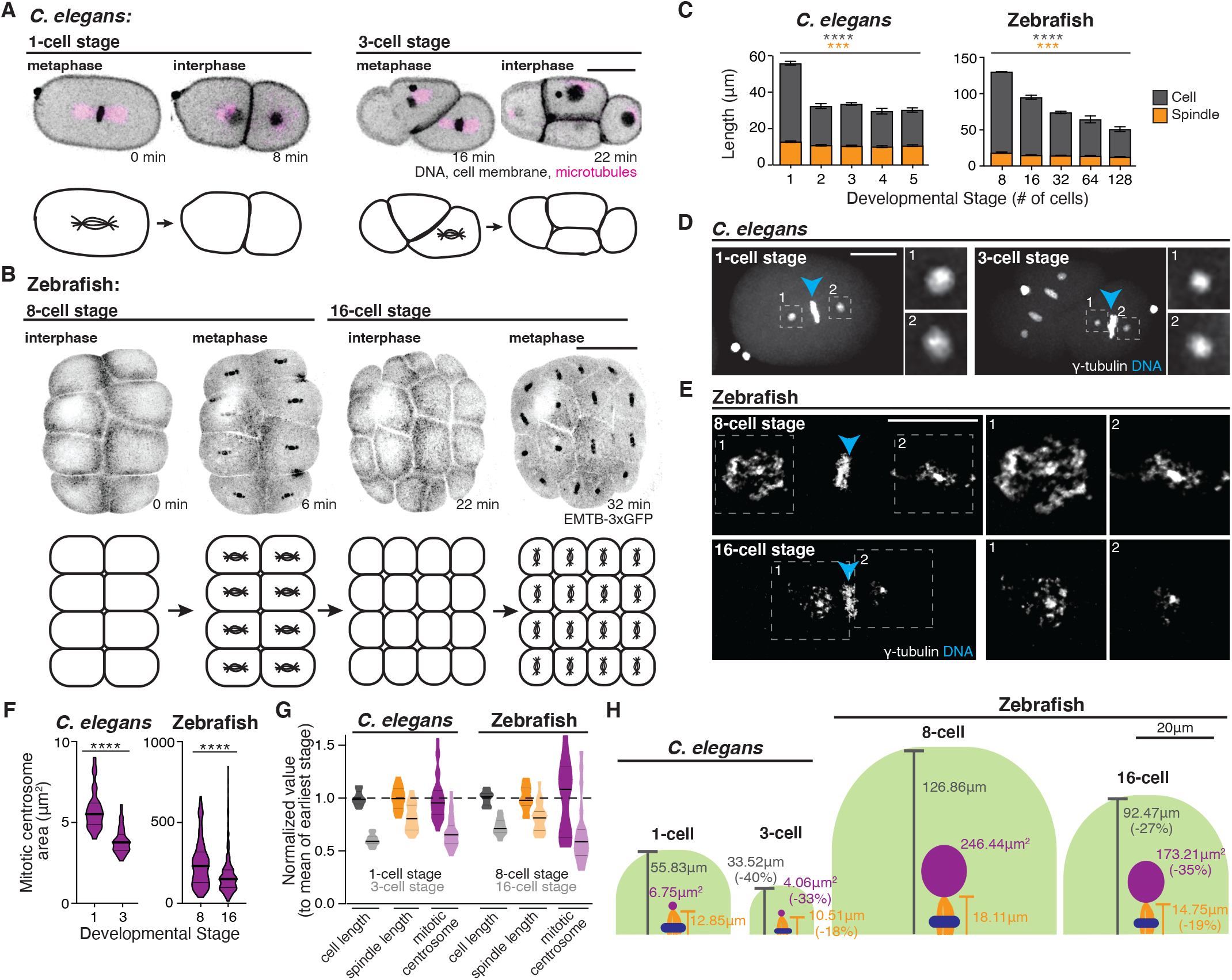
Mitotic centrosome area scales with cell length during embryonic cell divisions in *C. elegans* and zebrafish. **(A)** Maximum confocal projection of a *C. elegans* embryo at 1- and 3-cell stages. Nucleus and cell membrane (H2B∷mCherry and PH∷mCherry, inverted grayscale) and microtubules (α-tubulin∷GFP, magenta) shown. Model below depicting position of metaphase spindle within embryo. Bar, 10μm. **(B)** Three-dimensional rendering of zebrafish embryo at the 8- and 16-cell stage. Microtubule marker (EMTB-3xGFP) shown in grayscale. Model below depicting position of mitotic spindle within embryo. Bar, 250μm. **(C)** Bar graphs depicting spindle length (orange) and cell length along spindle axes (gray) during *C. elegans* (left, n=10 embryos) and zebrafish development (right, n=3 embryos). Mean ± SEM shown. One-way ANOVA, p<0.0001 (****) for *C. elegans* and zebrafish cell length, p=0.0005 (***) for *C. elegans* spindle length, p=0.0002 (***) for zebrafish spindle length. **(D-E)** Representative images of metaphase cell at the 1-cell (left) and 3-cell stage (right) in a *C. elegans* embryo (**D**), and at the 8-cell (top) and 16-cell stage (bottom) in a zebrafish embryo (**E**). Chromosomes and γ-tubulin shown in white, chromosomes denoted by blue arrowhead. Mitotic centrosomes highlighted in insets on right. Bar, 15μm. **(F)** Violin plot with box and whiskers depicting two-dimensional centrosome area (μm^2^) at the 1- and 3-cell stage in *C. elegans* (left, n>24 embryos), and at the 8- and 16-cell stage in zebrafish (right, n>12 embryos). Student’s t-test, p<0.0001 (****). **(G)** Violin plot depicting cell length (n>14, n>22), spindle length (n>14, n>22), and mitotic centrosome area for *C. elegans* at the 1- and 3-cell stage (n>46 centrosomes, left), and zebrafish at the 8- and 16-cell stage (n>147 centrosomes, right). Values normalized to mean of earliest developmental stage (1-cell for *C. elegans*, 8-cell for zebrafish), dashed line at value of 1. **(H)** Scaled model depicting cell (green), spindle (orange), and mitotic centrosome (purple) sizes during the 1- and 3-cell stage in *C. elegans* embryos, and the 8- and 16-cell stage in zebrafish embryos. Percentages listed at the 3-cell and 16-cell stage refer to the percent decrease in value compared to the previous developmental stage (rounded to the nearest percentage). Bar, 20μm. **For violin plots:** Plot boundaries depict minimum and maximum, 25^th^ and 75^th^ quartiles represented by thin black line, median represented by thick black line. **For all graphs:** detailed statistical analysis in Methods Tables.

In early development, rapid rounds of division result in a stark decrease in cell size during the cleavage stage [8]. We measured cell area during the first five cell cycles in each respective embryo (1- to 5-cell stage embryo in *C. elegans*, 8- to 128-cell stage in zebrafish). Since imaging and quantitative analysis is difficult to obtain from the 1-cell to 4-cell stage zebrafish embryo, we focused on the 8- to 128-cell stage. Both organisms had a significant decrease in cell area during these divisions. In *C. elegans*, the change in cell area became less drastic over time. In contrast, the decrease in cell area remained constant in zebrafish embryos out to 128-cells (Supplementary Figure 1E). This suggests that while a marked decrease in cell size occurs during the first several rounds of cell division in many organisms, the magnitude of this change is not always similar.

We next questioned whether the spindle and/or mitotic centrosome scaled to the longest cell axis (e.g. cell length) in *C. elegans* and zebrafish embryos. Spindle, mitotic centrosomes, and cell length were measured in *C. elegans* embryos that stably expressed a centrosome marker (*γ*-tubulin∷GFP), cell membrane marker (PH∷mCherry) and/or a nuclear marker (H2B∷mCherry) (Figure 1A, 1D, Supplementary Figure 1A-B, strains listed in methods). In zebrafish, live βactin:EMTB-3xGFP embryos or embryos fixed and stained for *γ*-tubulin were used to visualize microtubules (Figure 1B, Supplementary Figure 1A-B) or mitotic centrosomes (Figure 1E) respectively. Metaphase mitotic spindle length was measured from mitotic centrosome to mitotic centrosome, cell length was measured from cell membrane to cell membrane along the same plane of the metaphase spindle, and metaphase mitotic centrosome area was measured (modeled in Figure 1H). Cell length decreased with every division over time in *C. elegans* and zebrafish (Figure 1C, gray), similar to the trend identified in cell area (Supplementary Figure 1E). This decrease was not as drastic as the decrease in spindle length (Figure 1C, orange). When considered as a ratio between spindle and cell length, mitotic spindles occupy a higher percentage of the cell length in later cell divisions compared to earlier divisions in both organisms leading to a significant decrease in the distance from mitotic centrosomes to cell membrane (Supplementary Figure 1F-G). Despite the stark size difference between cells in *C. elegans* and zebrafish embryos (Figure 1H), these data suggest a conserved trend of disproportional changes in cell and spindle dimensions during early cell divisions.

When measuring mitotic centrosome size in *C. elegans* and zebrafish embryos (Figure 1D-E), a significant decrease in mitotic centrosome area was identified from one round of division to the next (Figure 1F). Strikingly, mitotic centrosomes in zebrafish embryos were extremely large with *γ*-tubulin organized into a wheel-like structure (246.44±11.93μm^2^ at 8-cell stage, Figure 1E), compared to *γ*-tubulin in *C. elegans* (6.75±0.28μm^2^ at 1-cell stage, Figure 1D) or in zebrafish at the 512-cell stage (3.16 ±1.36μm^2^, Supplementary Figure 1H).

The values obtained from cell length, spindle length, and mitotic centrosome area measurements were normalized to determine their relative change. Size values were normalized to the 1-cell stage in *C. elegans* and to the 8-cell stage in zebrafish. In both organisms, we determined that the change in cell length scaled more closely with the change in mitotic centrosome area than that of spindle length (Figure 1G). In both *C. elegans* and zebrafish embryos, cell length and mitotic centrosome area decreased by 30-40% over time. Spindle length, however, decreased <20% during this time in both organisms (Figure 1G). Taken together, these data suggest that decreases in cell size scale more closely with mitotic centrosome size than spindle length (Figure 1H, percentages represent size decrease).

### Centrosomes in early zebrafish development are uniquely structured

To characterize spindle and centrosome dynamics in the early embryo we focused on the zebrafish embryo due to its uniquely large centrosomes. To do this we employed βactin∷EMTB-3xGFP [7] and βactin∷centrin-GFP [9] embryos to mark microtubules and centrosomes. Volumetric projections of embryos from these transgenic lines were acquired over time (Figure 2A-B). The positioning of the mitotic spindles (Figure 2A) and mitotic centrosomes (Figure 2B, Supplementary Video 3) are consistent with that modeled in Figure 2C. At prophase, the mitotic centrosomes are placed on either side of the nucleus (Figure 2E, 2F) and begin to nucleate a robust microtubule-based spindle for metaphase (Figure 2D). During anaphase, the mitotic centrosomes begin to fragment and disperse, and reform during telophase to prepare for immediate cell cycle re-entry (Figure 2E-F, Supplementary Video 4).

**Figure 2.**
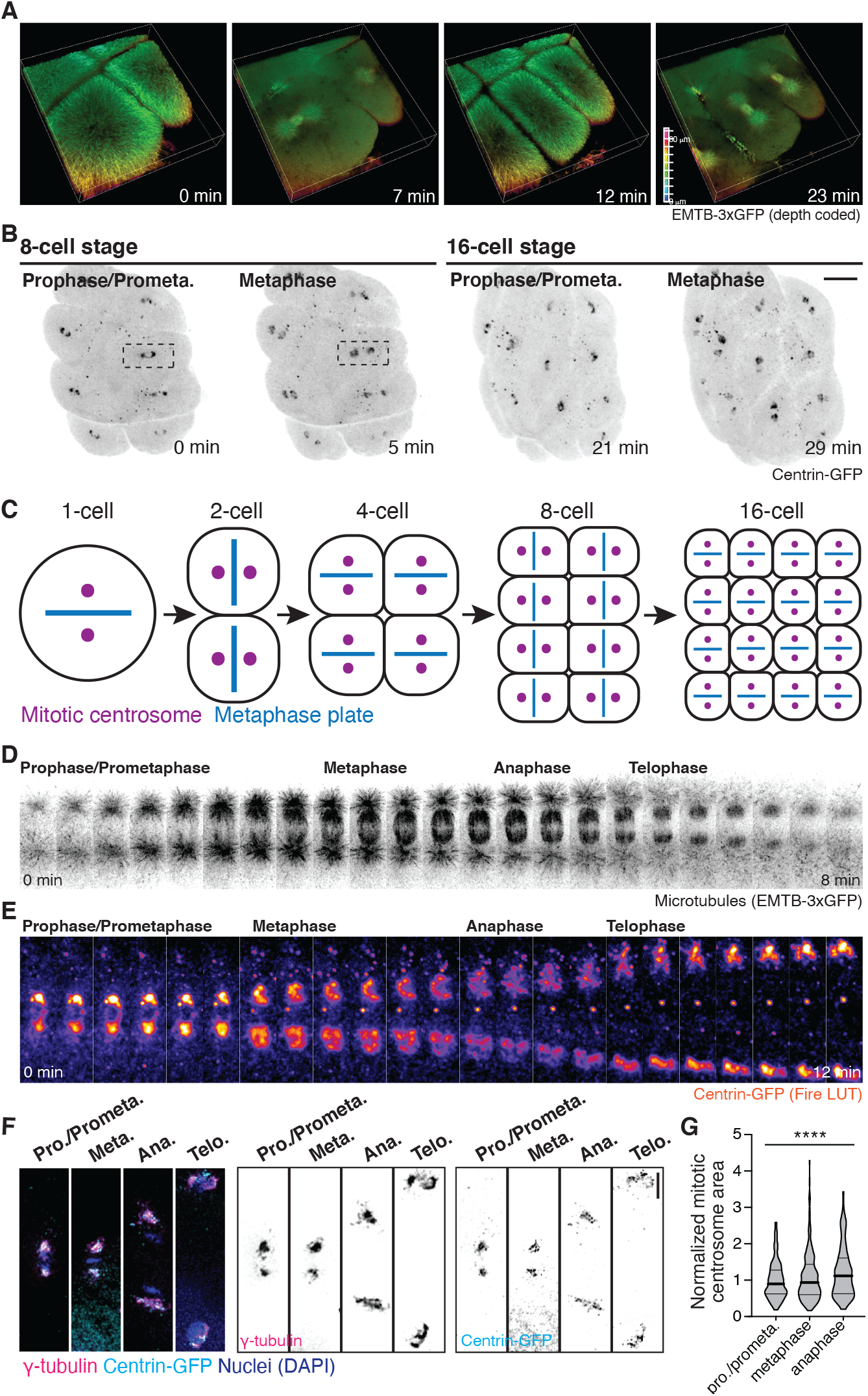
Centrosomes in early zebrafish development are uniquely structured. **(A-B)** Three-dimensional rendering of mitotic spindle positioning during early embryonic divisions in a zebrafish embryo using EMTB-3xGFP(microtubules, **A**) and centrin-GFP (centrosome, **B**). Microtubules shown in depth-coded z-stack such that z-slices closest to the embryo yolk are colored red and z-slices furthest from the yolk are colored blue (**A**). Centrin-GFP (inverted grayscale, **B**) shown at the 8- and 16-cell stage. Cell highlighted by dashed box magnified in (**E**). Bar, 100μm. **(C)** Model depicting the placement of mitotic spindles within embryonic zebrafish cells from the 1-cell stage to the 16-cell stage. Cells are viewed from top of cell mass with yolk placed below (XY view). Mitotic centrosomes (purple) and metaphase plate (blue) shown. Embryo midline placed perpendicular to spindle positioning drawn in 8- and 16-cell models. **(D-E)** Stills from timelapse of a cell division in EMTB-3xGFP transgenic embryo (microtubules, inverted grayscale, **D**) and a centrin-GFP embryo (centrosome, Fire LUT, **E**, insert from **B**). Mitotic stages denoted. **(F)** Single mitotic cells from fixed embryos in prometaphase, metaphase, anaphase, and telophase. Centrin-GFP (magenta/inverted grayscale), γ-tubulin (cyan/inverted grayscale), and nuclei (DAPI, blue) shown. Bar, 20μm **(G)** Violin plot depicting normalized mitotic centrosome area at 16-cell stage during prophase/prometaphase, metaphase, and anaphase. Values normalized to the mean mitotic centrosome area at prophase/prometaphase. One-way ANOVA, p<0.0001 (****). n>88 mitotic centrosomes measured. Plot boundaries depict minimum and maximum, 25^th^ and 75^th^ quartiles represented by thin black line, median represented by thick black line. Detailed statistical analysis in Methods Tables.

Notably, centrin normally marks centrioles [10,11], but in this case it marks a uniquely large structure that colocalizes with the PCM protein *γ*-tubulin [12] (Figure 2F, Supplemental Figure 2A-B). When measuring *γ*-tubulin area, the mitotic centrosome significantly increased in size between prophase and anaphase (Figure 2F, 2G), with centrin maintaining colocalization at these mitotic stages (Figure 2F, Supplementary Figure 2B). This points to a unique centrosome structure that is specific to the extremely large zebrafish embryo cells. The degree in colocalization increases when comparing 8-cell, 16-cell, and 512-cell stage embryos (Supplementary Figure 2B) suggesting that centrin and *γ*-tubulin distribution at the centrosome changes during zebrafish development.

### Mitotic centrosomes are asymmetric during early zebrafish cell divisions

An asymmetry in mitotic centrosome size was identified during early zebrafish embryonic divisions with the largest metaphase mitotic centrosome pointing towards the midline of the embryo cell grid (refer to dashed orange line to mark embryo midline in Figure 3A) that wasn’t observed in *C. elegans* embryos (Figure 1D). A greater than 2-fold difference was calculated when comparing the measured area between the larger and smaller mitotic centrosome and the inner and outer mitotic centrosome at the 8-cell and 16-cell stage (Figure 3B, Supplementary Figure 3A, 3C) with the largest mitotic centrosome consistently orienting towards the embryo midline (88.5± 2.7 % of the time across n=36 embryos at the 8- and 16-cell stage measured, Figure 3A). This > 2-fold difference in mitotic centrosome size is maintained from prophase/prometaphase to anaphase (Supplementary Figure 3B, D). This asymmetry in mitotic centrosome size was consistent in cells placed next to the midline or further away from the midline (representative shown in Figure 3A).

**Figure 3.**
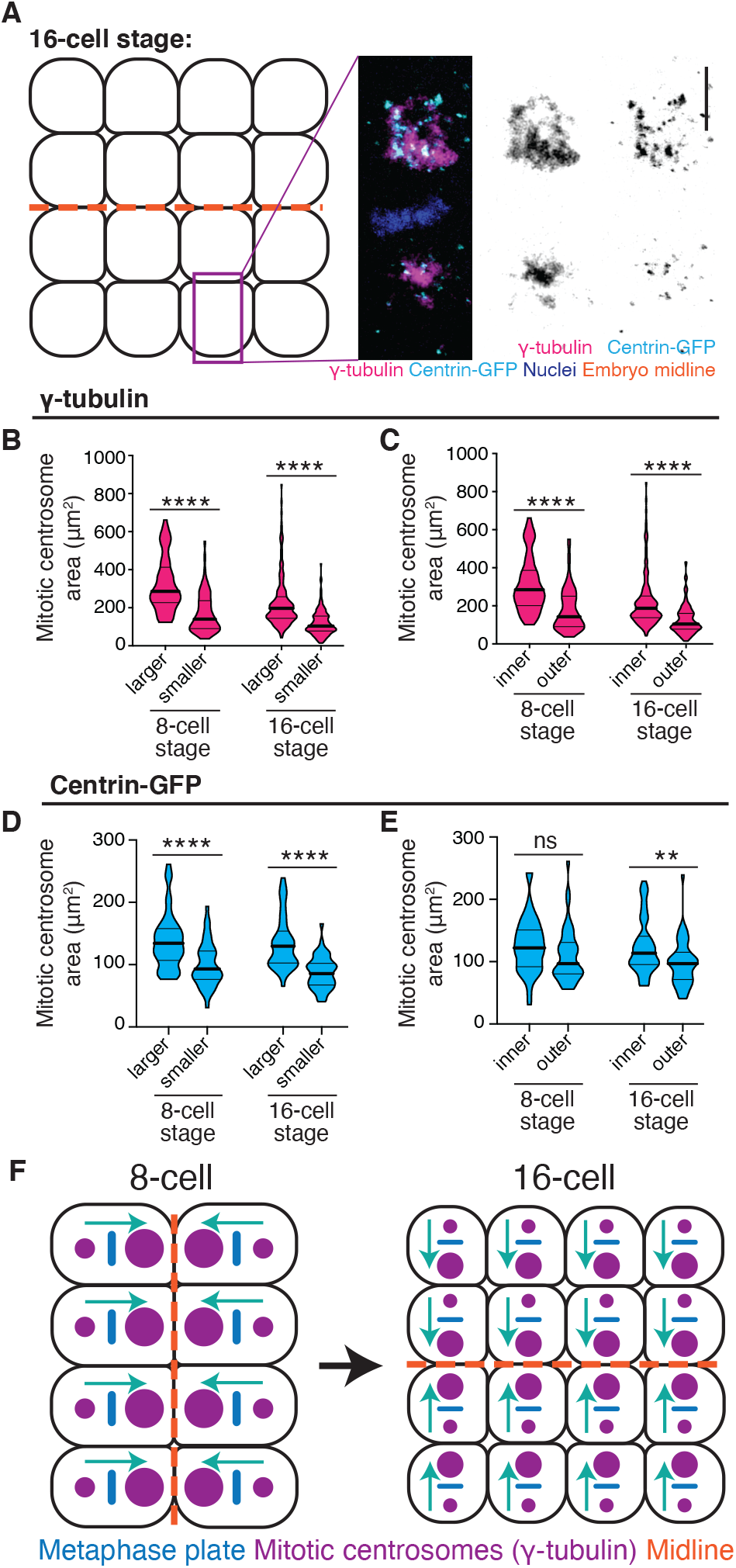
Mitotic centrosomes are asymmetric during early zebrafish cell divisions. **(A)** Model depicting a 16-cell embryo with maximum confocal projections of a representative cell (denoted with purple box). Fixed 16-cell metaphase embryo expressing centrin-GFP (magenta/inverted grayscale) and immunostained for γ-tubulin (cyan/inverted grayscale) and nuclei (DAPI, blue). Embryonic midline denoted with orange dashed line. Bar, 10μm. **(B-E)** Violin plot depicting the mitotic centrosome area in γ-tubulin-labeled embryos (**B-C**) or centrin-GFP embryos (**D-E**) at the 8- and 16-cell stage binned by size (larger/smaller in **B, D**) or position relative to midline (inner/outer, **C, E**). Students t-test, p<0.0001 (****, **B-C**). N>13 embryos, p<0.0001 (****, **D**), p=0.1183 (ns, **E**), and p=0.0030 (**, **E**). N>6 embryos (**D-E**). Plot boundaries depict minimum and maximum, 25^th^ and 75^th^ quartiles represented by thin black line, median represented by thick black line. **(F)** Model depicting the positioning of the asymmetric mitotic centrosomes in relation to the embryonic midline during the 8- and 16-cell stages. The larger of the two mitotic centrosomes (purple) is placed closest to the embryonic midline (orange), providing directionality (turquoise arrow). **For all graphs:** detailed statistical analysis in Methods Tables.

When calculating centrosome size at metaphase in the centrosome transgenic line, βactin∷centrin-GFP, an approximate 1.5-fold change was calculated at the 8-cell and 16-cell stage (Figure 3D, Supplementary Figure 3E). Even though centrin-GFP organization at mitotic centrosomes is asymmetric (Figure 3D), a metaphase spindle with the mitotic centrosome organizing the largest area of centrin-GFP was not as consistently positioned towards the center of the embryo (60.1± 4.8 % of the time across n=16 embryos at the 8- and 16-cell stage measured, Figure 3E, ratios between mitotic centrosomes shown for inner to outer in Supplemental Figure 3F).

Taken together, these data suggest a model in which zebrafish mitotic centrosomes present with an asymmetry in PCM components (e.g. γ-tubulin) starting at prophase/prometaphase (Supplementary Figure 3B, 3D) and this asymmetry biases the positioning of the larger centrosome towards the midline at the 8- and 16-cell stage (modeled in Figure 3F).

### PLK1 and PLK4 activity are required for asymmetric mitotic centrosome positioning

As cells progress through the cell cycle, they normally require PLK4 to duplicate their centrosome and PLK1 for robust PCM assembly during bipolar spindle construction [13]. The assembly of PCM components that interact with *γ*-tubulin, such as pericentrin and CEP215, is facilitated by the phosphorylation activity of PLK1 [13,14]. With PLK4 inhibition, centriole duplication is disrupted, causing spindles to assemble through acentriolar organization of PCM [15–17]. However, the role of PLK1 and/or PLK4 at mitotic centrosomes in the early zebrafish embryo is unknown. Transcripts for PLK1 and PLK4 have been detected as early as the 1-cell stage in zebrafish embryos, indicating that they are maternally supplied prior to zygotic genome activation, albeit PLK4 transcript levels are significantly lower [18]. Due to this, we tested the hypothesis that PLK1 and/or PLK4 regulate *γ*-tubulin organization at mitotic centrosomes in zebrafish embryos. The PLK1 and PLK4 small molecule inhibitors, BI2536 [10,19,20] and centrinone [21], were injected into 1 cell stage embryos (concentrations used 1μM or 100nM, BI2536 described with zebrafish in [19,20]). An injection approach was utilized versus soaking the embryos in drug due to the low permeability of the zebrafish chorion and embryo fragility at this stage. Control embryos were injected with 1% DMSO at the 1-cell stage and analyzed at the 16-cell stage. In 87.14±4.16% of these control embryos, the larger mitotic centrosome was positioned towards the midline (101.65±4.91μm^2^) whereas the smaller was positioned away (52.28±272μm^2^, Figure 4A-B). This directional positioning of the larger mitotic centrosome towards the midline was significantly decreased when embryos were injected with BI2536 (60.92±5.11% with 100nM and 44.78±7.18% with 1μM), or centrinone (66.13±8.17%, with 100nM and 60.67±6.87% with 1μM, Figure 4A-B). Interestingly, the ratio of mitotic centrosome size difference within a spindle decreased and the overall centrosome area increased in embryos injected with BI2536 or centrinone (Figure 4C-D, Supplemental Figure 4A-C). This suggests that not only does PLK1 and PLK4 regulate mitotic centrosome structure and asymmetry, but they regulate the directionality of larger centrosome placement towards the midline of the embryo’s grid of cells (Figure 4E).

**Figure 4.**
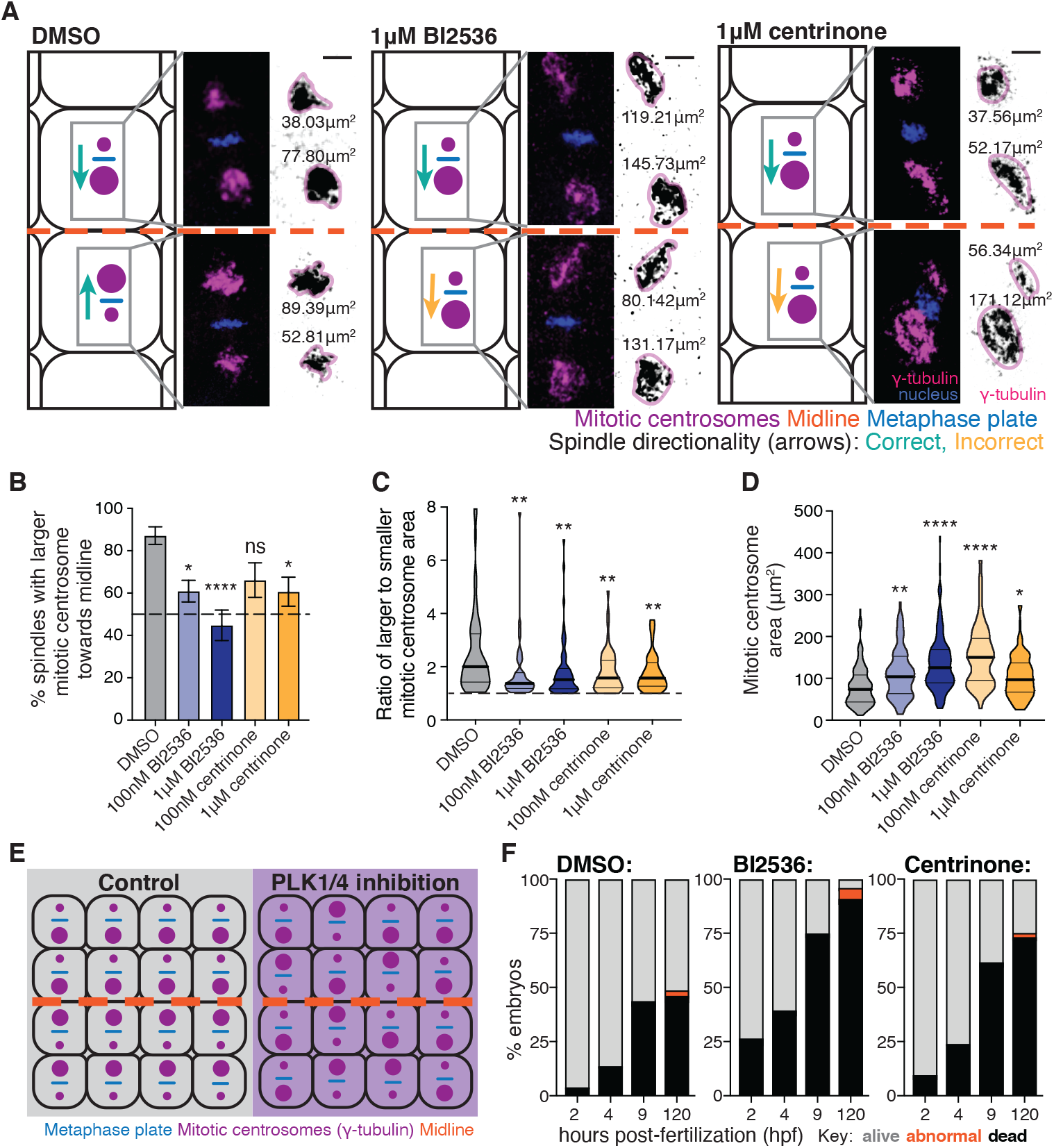
PLK1 and PLK4 activity are required for asymmetric mitotic centrosome positioning. **(A)** Representative images of 16-cell stage embryos during metaphase under conditions of DMSO (left), 1μM BI2536 (center), or 1μM centrinone treatment (right). Single cells denoted in embryo image magnified in inset. γ-tubulin (magenta/inverted grayscale), and nuclei (DAPI, blue) shown. Model depicting mitotic centrosome positioning in embryo shown on left (cyan/correct and gold/incorrect positioning depicted with arrows). Large and smaller mitotic centrosomes not drawn to scale in model. Bar, 100μm. **(B)** Bar graph depicting percentage of spindles with largest centrosome pointed towards midline under conditions of DMSO (gray), BI2536 (100nM or 1μM, blue), or centrinone (100nM or 1μM, gold) exposure. **(C)** Violin plot depicting the ratio of mitotic centrosome areas binned by size (larger-to-smaller centrosome ratio) under conditions of DMSO (gray), BI2536 (100nM or 1μM, blue), or centrinone (100nM or 1μM, gold) exposure. Mitotic centrosome areas measured from γ-tubulin signal from fixed zebrafish embryos at the 16-cell stage. **(D)** Violin plot depicting mitotic centrosome area measured from γ-tubulin signal from fixed zebrafish embryos at the 16-cell stage under conditions of DMSO (gray), BI2536 (100nM or 1μM, blue), or centrinone (100nM or 1μM, gold) exposure. One-way ANOVA with Dunnett’s multiple comparison test performed with DMSO control. **(E)** Model depicting the positioning of the asymmetric mitotic centrosomes in relation to the embryonic midline during the 16-cell stages under conditions of DMSO (gray), or BI2536 or centrinone (purple) exposure. Mitotic centrosomes (purple), metaphase plate (blue), and embryonic midline (orange dashed line) depicted. **(F)** Bar graphs representing percentage of embryos alive (gray), dead (black), or with abnormal phenotypes (orange) at 2, 4, 9, and 120 hours post-fertilization (hpf) after treatment with 1% DMSO vehicle control (left, n=80 embryos), 1μM BI2536 (center, n=200 embryos), or 1μM centrinone (right, n=154 embryos). **For all graphs:** detailed statistical analysis in Methods Tables.

We were surprised that PLK1 inhibition caused an increase in the area occupied by *γ*-tubulin in mitotic centrosomes due to its known role in recruiting the pericentrin-CEP215 complex that anchors the γ-TURC at the centrosome [13,22]. One possible explanation for this is that PLK1 has been proposed to regulate PCM architecture by facilitating its phase separation in *C. elegans* [23,24] and that inhibiting PLK1 in zebrafish embryos may change the physical state of the PCM causing it to increase in size. This could explain why mitotic centrosome area significantly increases in a dosage-dependent manner with BI2536 treatment, as losing a PCM architecture regulator may cause the surrounding PCM to lose its tight matrix configuration and expand in size (Figure 4A, D). Centrinone treatment did not exhibit the same dosage-dependent change in mitotic centrosome area (Figure 4D). A possible explanation is that PLK4 is present at much lower concentrations in the early zebrafish embryo compared to PLK1 [18]. It is therefore likely that lower drug concentrations are required to target the small pool of PLK4, leading to a similar phenotype with drug concentrations above this small threshold.

In order to determine the importance of PLK1/4-dependent asymmetric mitotic centrosome size placement in early zebrafish divisions, we raised embryos after injection of 1% DMSO, 1μM BI2536, or 1μM centrinone (Figure 4F). We found that compared to control embryos, PLK1- or PLK4-inhibition resulted in a lower survival rate over the first five days post-fertilization. At five days, we noted heart edema, embryo elongation defects, yolk elongation defects, and small eyes in the small fraction of embryos that survived drug treatment (Figure 4F, Supplementary Figure 4D), which are all common defects seen after early development perturbation. Given that the injections of BI2536 or centrinone likely diffuse out when the chorion starts to become more permeable, it is probable that the earliest cell divisions are impacted the most from this treatment. This led us to conclude that PLK1- and PLK4-dependent asymmetric mitotic centrosome placement in early embryos impacts later development.

Through these studies, a unique centrosome structure has been characterized that may contribute to a better understanding of how mitotic spindles are able to coordinate cell division in disproportionately large cells. We demonstrate that mitotic centrosome size adapts to the decreasing cell size during the cleavage stage of zebrafish development. During this time in development, we found that zebrafish mitotic centrosomes are asymmetric in size and display a directionality, where the larger mitotic centrosome within a spindle is positioned towards the embryonic midline in a PLK1- and PLK4-dependent manner. Furthermore, we identified an ability for mitotic centrosomes to scale with cell size and maintain a 2-fold asymmetry across the spindle while doing so. When centrosome size scaling and asymmetry is disrupted an increase in embryonic lethality and developmental defects occurs, suggesting that early zebrafish embryonic cell divisions are not only important for early embryogenesis but likely also impact later developmental processes.

## ACKNOWLEDGEMENTS

We thank Lilianna Solnica-Krezel (UW) for sharing the βactin∷centrin-GFP zebrafish line. This work was supported by National Institutes of Health Grants no. R00GM107355, no. R01GM127621 (to H.H.), no. R01 GM114471 (to J.N.B), and a Syracuse University ‘Cuse Good-to-Great award. We thank the *Caenorhabditis* Genetics Center, which is funded by National Institutes of Health Office of Research Infrastructure Programs (P40OD010440), for providing strains for this study.

## Methods

### Animal Lines

Zebrafish lines were maintained using standard procedures approved by the Syracuse University IACUC committee (protocol #18-006). Embryos were staged as described in Kimmel et al 1995. Wildtype zebrafish lines as well as transgenic lines were used for live imaging and immunohistochemistry. Transgenic *C.* elegans lines were imaged and characterized by the Bembenek lab. See Methods Tables for list of transgenic zebrafish and *C. elegans* lines used.

### Imaging

For zebrafish, a Leica SP5 or SP8 (Leica, Bannockburn, IL) laser scanning confocal microscope (LSCM) was used throughout manuscript. A HC PL APO 20x/0.75 IMM CORR CS2 objective, HC PL APO 40x/1.10 W CORR CS2 0.65 water immersion objective, and an HCX Plan Apochromat 63×/1.40-0.06 NA OIL objective were used. Images were acquired using LAS-X software. Images taken with the SP8 LSCM were obtained through lightning, a built-in deconvolution algorithm. A Leica DMi8 (Leica, Bannockburn, IL) with a X-light v2 confocal unit spinning disk was also used, equipped with an 89 North - LDI laser and a Photometrics Prime-95B camera. Optics used were either 10x/0.32 NA air objective, HC PL APO 63X/1.40 NA oil CS2, HC PL APO 40X/1.10 NA WCS2 CORR, a 40X/1.15 N.A. 19 Lamda S LWD, or 100Å~/1.4 N.A. HC Pl Apo oil emersion objective. A Leica M165 FC stereomicroscope equipped with DFC9000 GT sCMOS camera was used for phenotypic analysis of embryos.

For live cell imaging of *C. elegans* embryos, a spinning disk confocal system was used. The system is equipped with a Nikon Eclipse and is an inverted microscope with a 60X 1.40NA objective, a CSU-22 spinning disc system and a Photometrics EM-CCD camera from Visitech International. Images were obtained every 2 minutes with a 1-micron z-stack step size.

### Pharmacological treatments

Zebrafish embryos were injected with either 1% DMSO, or BI2536 or centrinone (final concentration 100nM or 1μM) at the 1-2-cell stage. Embryos are incubated at 30°C until they reach the developmental stage of interest, at which time they are fixed with 4% paraformaldehyde in PBS. Immunohistochemistry then proceeds as detailed below.

### Zebrafish immunohistochemistry

Zebrafish embryos were fixed using 4% PFA containing 0.5% Triton-X 100 overnight at 4°C. Zebrafish were then dechorionated and incubated in PBST (phosphate buffered saline + 0.1% Tween) for 30 minutes. Embryos were blocked using a Fish Wash Buffer (PBS + 1% BSA + 1% DMSO + 0.1% Triton-X 100) for 30 minutes followed by primary antibodies incubation (antibodies diluted 1:200 in Fish Wash Buffer) overnight at 4°C or 3 hours at room temperature. Embryos are then washed five times in Fish Wash Buffer and incubated in secondary antibodies (diluted 1:200 in Fish Wash Buffer) for 3 hours at room temperature. After five more washes, embryos were incubated with 4’,6-diamidino-2-phenylindole (NucBlue® Fixed Cell ReadyProbes® Reagent) for 30 minutes. For imaging, embryos were either halved and mounted on slides using Prolong Diamond (Thermo Fisher Scientific cat. # P36971) or whole-mounted in 2% agar (Thermo-Fisher cat. # 16520100).

### Image and Data Analysis

Images were processed using both FIJI/ImageJ software and Adobe Photoshop. All graphs and statistical analysis were produced using Graphpad Prism software. 3-D images, movies, and surface rendering were performed using Bitplane IMARIS software (Surface, Smoothing, Masking, and Thresholding functions).

To calculate two-dimensional area, a boundary was drawn around the structure of interest (cell, spindle pole, etc.) in ImageJ/FIJI and the area within this shape was calculated. To calculate spindle length, cell length, aspect ratio, etc., a line was drawn in ImageJ/FIJI from one end of the structure of interest to the other. This length was then measured and recorded.

### Phenotypic characterization

Wildtype zebrafish embryos were injected as described in the pharmacological section described above. The embryos were maintained at 30°C and assessed for abnormality in development and the number of deaths every 30 minutes for 9-10 hours post injection then once 24 hours post injection. At 5 days post fertilization, the phenotypes of injected embryos were characterized and the number of embryos with developmental defects were recorded.

To generate death curves for the pharmacological treatments, the number of embryos treated with each drug were standardized to the starting number of embryos and were displayed as ratios over time.

### Statistical analysis

Unpaired, two-tailed Student’s t-tests and one-way ANOVA analyses were performed using GraphPad Prism software. **** depicts a p-value <0.0001, *** p-value <0.001, **p-value<0.01, *p-value <0.05. See Methods Tables for detailed information regarding statistics.

## KEY RESOURCES TABLE

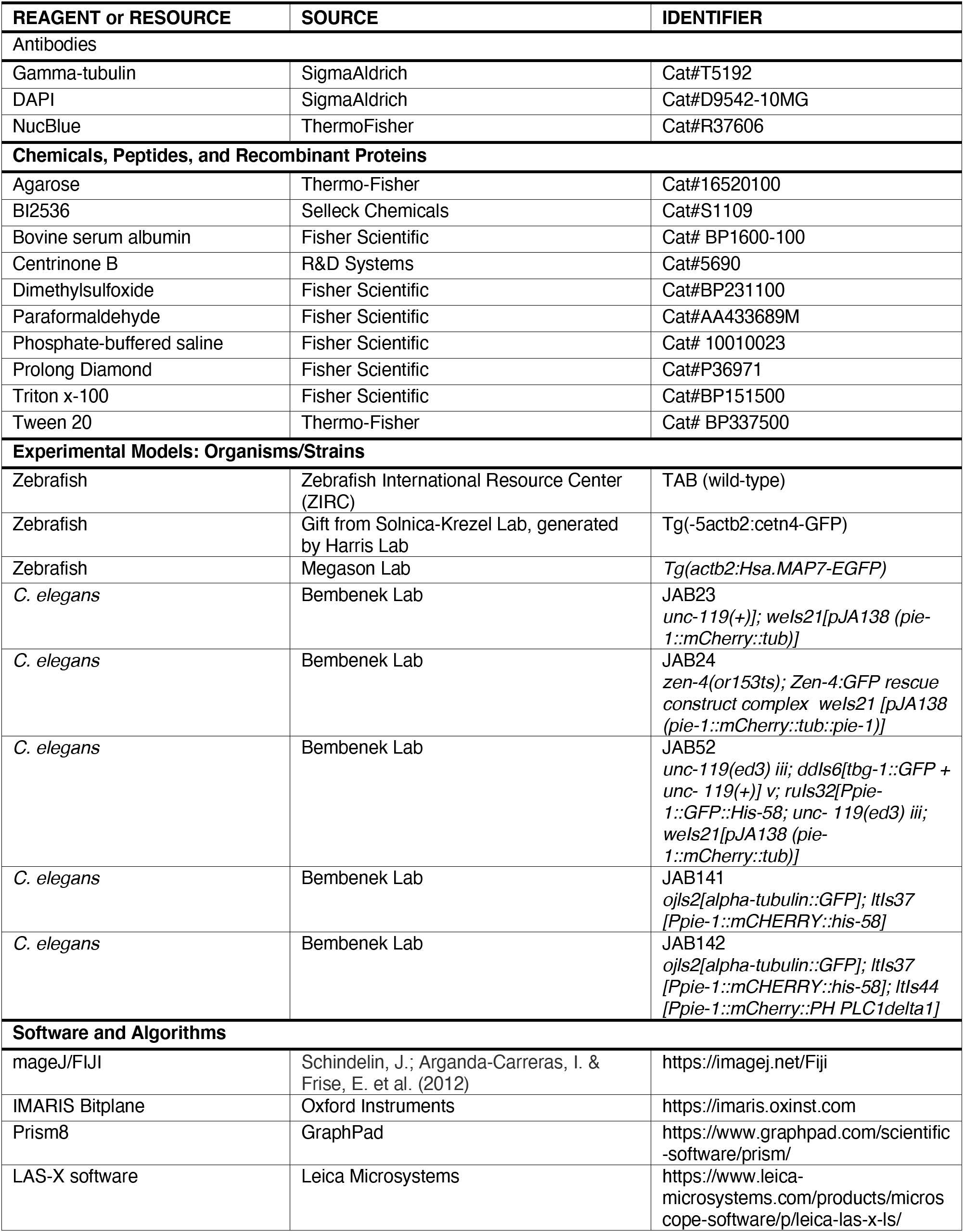

## Methods Table: Detailed statistical analysis results

**Figure 1:**
| Figure | Category | n | Statistical test | Parameters | Result | p-value |
| --- | --- | --- | --- | --- | --- | --- |
| 1C | *C. elegans* cell | 10 | One-way ANOVA | F (4, 69) = 88.18 | **** | <0.0001 |
|  | *C. elegans* spindle | 10 |  | F (4, 81) = 5.558 | *** | 0.0005 |
|  | Zebrafish cell | 3 |  | F (4, 10) = 109.6 | **** | <0.0001 |
|  | Zebrafish spindle | 3 |  | F (4, 10) = 16.69 | *** | 0.0002 |
| 1F | *C. elegans* | 13 | Two-tailed Student's t-test | t=8.975, df=92 | **** | <0.0001 |
|  | Zebrafish | 147, 346 |  | t=5.832, df=491 | **** | <0.0001 |
| S1E | *C. elegans* | 10 | One-way ANOVA | F (4, 45) = 611.6 | **** | <0.0001 |
|  | Zebrafish | 3 |  | F (4, 10) = 105.5 | **** | <0.0001 |
| S1F | *C. elegans* | 10 | One-way ANOVA | F (4, 69) = 16.57 | **** | <0.0001 |
|  | Zebrafish | 3 |  | F (4, 10) = 26.35 | **** | <0.0001 |
| S1G | *C. elegans* | 10 | One-way ANOVA | F (9, 308) = 83.44 | **** | <0.0001 |
|  | Zebrafish | 3 |  | F (4, 10) = 114.8 | **** | <0.0001 |

**Figure 2:**
| Figure | Category | n | Statistical test | Parameters | Result | p-value |
| --- | --- | --- | --- | --- | --- | --- |
| 2G | Prophase/prometaphase | 90 | One-way ANOVA | F (2, 389) = 10.62 | **** | <0.0001 |
|  | Metaphase | 214 |  |  |  |  |
|  | Anaphase | 88 |  |  |  |  |
| S2B | 8-cell | 23 | One-way ANOVA | F (2, 53) = 3.198 | * | 0.0488 |
|  | 16-cell | 19 |  |  |  |  |
|  | 512-cell | 14 |  |  |  |  |

**Figure 3:**
| Figure | Category | n | Statistical test | Parameters | Result | p-value |
| --- | --- | --- | --- | --- | --- | --- |
| 3B | 8-cell | 73 | Two-tailed Student's t-test | t=7.258, df=144 | **** | <0.0001 |
|  | 16-cell | 172 |  | t=9.509, df=342 | **** | <0.0001 |
| 3C | 8-cell | 73 | Two-tailed Student's t-test | t=6.231, df=145 | **** | <0.0001 |
|  | 16-cell | 172 |  | t=8.119, df=332 | **** | <0.0001 |
| 3D | 8-cell | 41 | Two-tailed Student's t-test | t=4.337, df=80 | **** | <0.0001 |
|  | 16-cell | 45 |  | t=6.406, df=88 | **** | <0.0001 |
| 3E | 8-cell | 41 | Two-tailed Student's t-test | t=1.579, df=80 | ns | 0.1183 |
|  | 16-cell | 45 |  | t=3.052, df=88 | ** | 0.003 |
| S3A | 8-cell | 73 | Two-tailed Student's t-test | t=0.3721, df=238 | ns | 0.7101 |
|  | 16-cell | 172 |  |  |  |  |
| S3B | Prophase/prometaphase | 45 | Two-tailed Student's t-test | t=6.225, df=88 | **** | <0.0001 |
|  | Metaphase | 107 |  | t=8.973, df=212 | **** | <0.0001 |
|  | Anaphase | 44 |  | t=5.713, df=86 | **** | <0.0001 |
| S3C | 8-cell | 73 | Two-tailed Student's t-test | t=0.3561, df=238 | ns | 0.7221 |
|  | 16-cell | 172 |  |  |  |  |
| S3D | Prophase/prometaphase | 45 | Two-tailed Student's t-test | t=4.678, df=88 | **** | <0.0001 |
|  | Metaphase | 107 |  | t=6.026, df=212 | **** | <0.0001 |
|  | Anaphase | 44 |  | t=4.987, df=86 | **** | <0.0001 |
| S3E | 8-cell | 41 | Two-tailed Student's t-test | t=1.287, df=84 | ns | 0.2015 |
|  | 16-cell | 45 |  |  |  |  |
| S3F | 8-cell | 41 | Two-tailed Student's t-test | t=1.274, df=84 | ns | 0.2063 |
|  | 16-cell | 45 |  |  |  |  |

**Figure 4:**
| Figure | Category | n | Statistical test | Parameters | Result | p-value |
| --- | --- | --- | --- | --- | --- | --- |
| 4B | DMSO | 13 | One-way ANOVA with Dunnett's multiple comparison | $F(5, 54) = 5.594$ | control | control |
|  | 100nM BI2536 | 12 |  |  | ns | 0.0312 |
|  | 1uM BI2536 | 13 |  |  | *** | <0.0001 |
|  | 100nM centrinone | 9 |  |  | ns | 0.1178 |
|  | 1uM centrinone | 7 |  |  | * | 0.0239 |
| 4C | DMSO | 107 | One-way ANOVA with Dunnett's multiple comparison | $F(5, 279) = 4.129$ | control | control |
|  | 100nM BI2536 | 93 |  |  | ** | 0.0019 |
|  | 1uM BI2536 | 128 |  |  | **** | 0.0059 |
|  | 100nM centrinone | 52 |  |  | * | 0.0055 |
|  | 1uM centrinone | 28 |  |  | * | 0.0054 |
| 4D | DMSO | 214 | One-way ANOVA with Dunnett's multiple comparison | $F(5, 564) = 24.61$ | control | control |
|  | 100nM BI2536 | 186 |  |  | ns | 0.0013 |
|  | 1uM BI2536 | 264 |  |  | *** | <0.0001 |
|  | 100nM centrinone | 66 |  |  | **** | <0.0001 |
|  | 1uM centrinone | 56 |  |  | ** | 0.0215 |
| S4A | DMSO | 107 | Two-tailed Student's t-test | $t=7.732, df=116$ | **** | <0.0001 |
| | 100nM BI2536 | 93 | | $t=4.110, df=94$ | **** | <0.0001 |
| | 1uM BI2536 | 128 | | $t=6.159, df=118$ | **** | <0.0001 |
| | 100nM centrinone | 52 | | $t=6.437, df=108$ | **** | <0.0001 |
| | 1uM centrinone | 28 | | $t=5.513, df=94$ | **** | <0.0001 |
| S4B | DMSO | 107 | Two-tailed Student's t-test | $t=6.441, df=116$ | control | control |
| | 100nM BI2536 | 93 | | $t=0.9344, df=94$ | ns | 0.3525 |
| | 1uM BI2536 | 128 | | $t=0.6171, df=118$ | ns | 0.5383 |
| | 100nM centrinone | 52 | | $t=2.095, df=108$ | * | 0.0385 |
| | 1uM centrinone | 28 | | $t=1.553, df=94$ | ns | 0.1238 |
| S4C | DMSO | 107 | One-way ANOVA with Dunnett's multiple comparison | $F(5, 279) = 7.769$ | control | control |
|  | 100nM BI2536 | 93 |  |  | *** | 0.0003 |
|  | 1uM BI2536 | 128 |  |  | **** | <0.0001 |
|  | 100nM centrinone | 52 |  |  | *** | 0.0001 |
|  | 1uM centrinone | 28 |  |  | *** | 0.0002 |

---

**Supplementary Figure 1.**
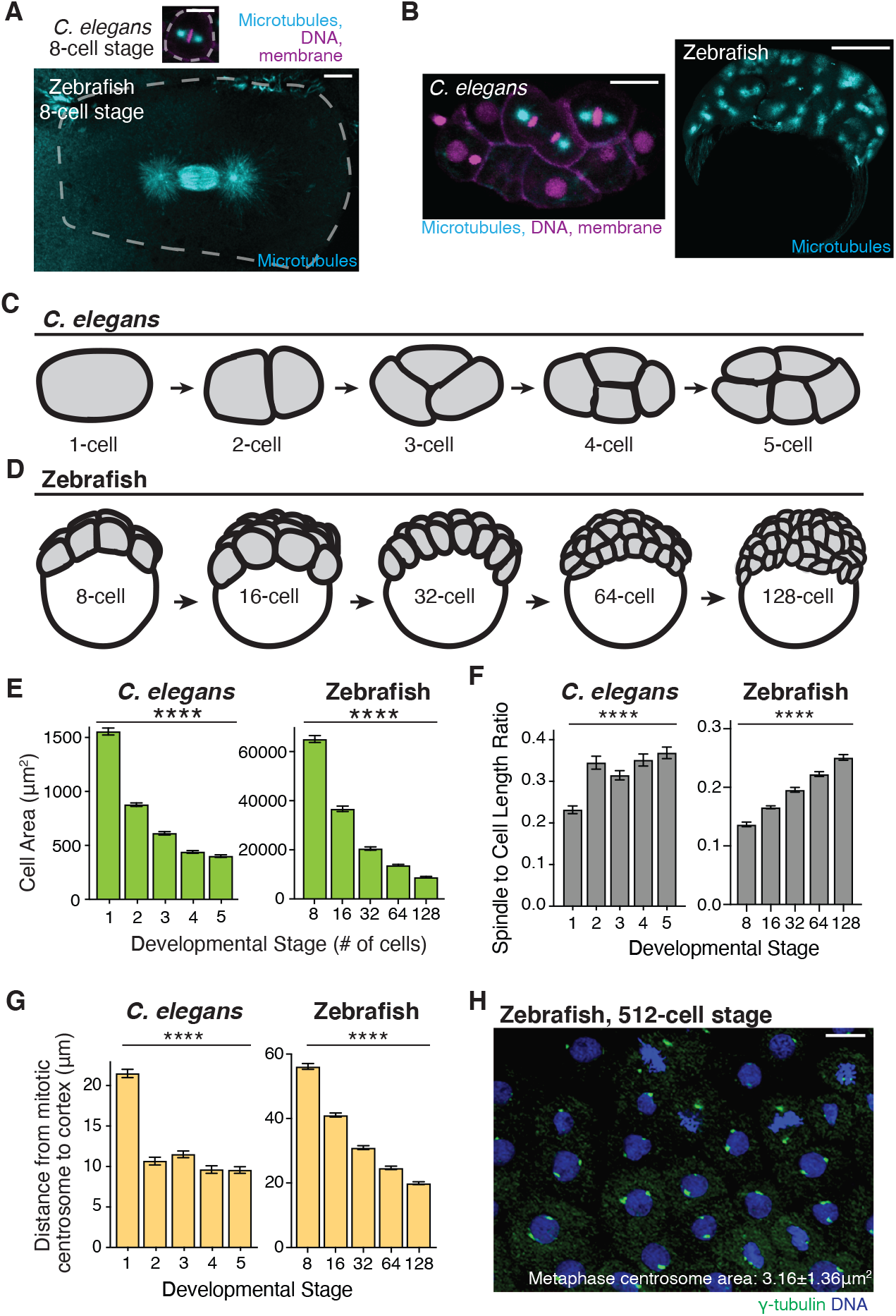
Mitotic centrosome area scales with cell length during embryonic cell divisions in *C. elegans* and zebrafish. **(A)** Representative images of a single cell at the 8-cell stage in a *C. elegans* embryo (left) and zebrafish embryo (right). Microtubules (cyan), cell membrane and histone marker (magenta) shown with cell boundary highlighted in white. Bar, 10μm. **(B)** Representative images of *C. elegans* (left) and zebrafish embryos (right) during the first few hours after fertilization. Microtubules (cyan), cell membrane and histone marker (magenta) shown. Bar, 20μm and 200μm, respectively. **(C)** Model of early *C. elegans* developmental stages (1-through 5-cell stage). Embryonic cells shown in gray. **(D)** Model of early zebrafish developmental stages (8-through 128-cell stage). Embryonic cells shown in gray, yolk shown in white. **(E)** Bar graph depicting two-dimensional cell area of single cells during *C. elegans* (left, n=10 embryos) and zebrafish embryo development (right, n=3 embryos). Mean ± SEM shown. One-way ANOVA, p<0.0001 (****). **(F)** Bar graph depicting the ratio of spindle length to cell length in *C. elegans* (n=10 embryos) and zebrafish embryos (n=3 embryos). Values were calculated by dividing the spindle length (Figure 1C, orange bars) by the cell length (1C, gray bars) to determine the percentage of the cell length occupied by the spindle. Mean ± SEM shown. One-way ANOVA, p<0.0001 (****) for both *C. elegans* and zebrafish. **(G)** Bar graph depicting the distance from mitotic centrosome to cell cortex in *C. elegans* (n=10 embryos) and zebrafish embryos (n=3 embryos) during early development. Mean ± SEM shown. One-way ANOVA, p<0.0001 (****) for both *C. elegans* and zebrafish. **(H)** Representative image of mitotic centrosome morphology at the 512-cell stage of zebrafish development. γ-tubulin (green) and chromosomes (blue) shown. **For all graphs:** detailed statistical analysis in Methods Tables.

**Supplementary Figure 2.**
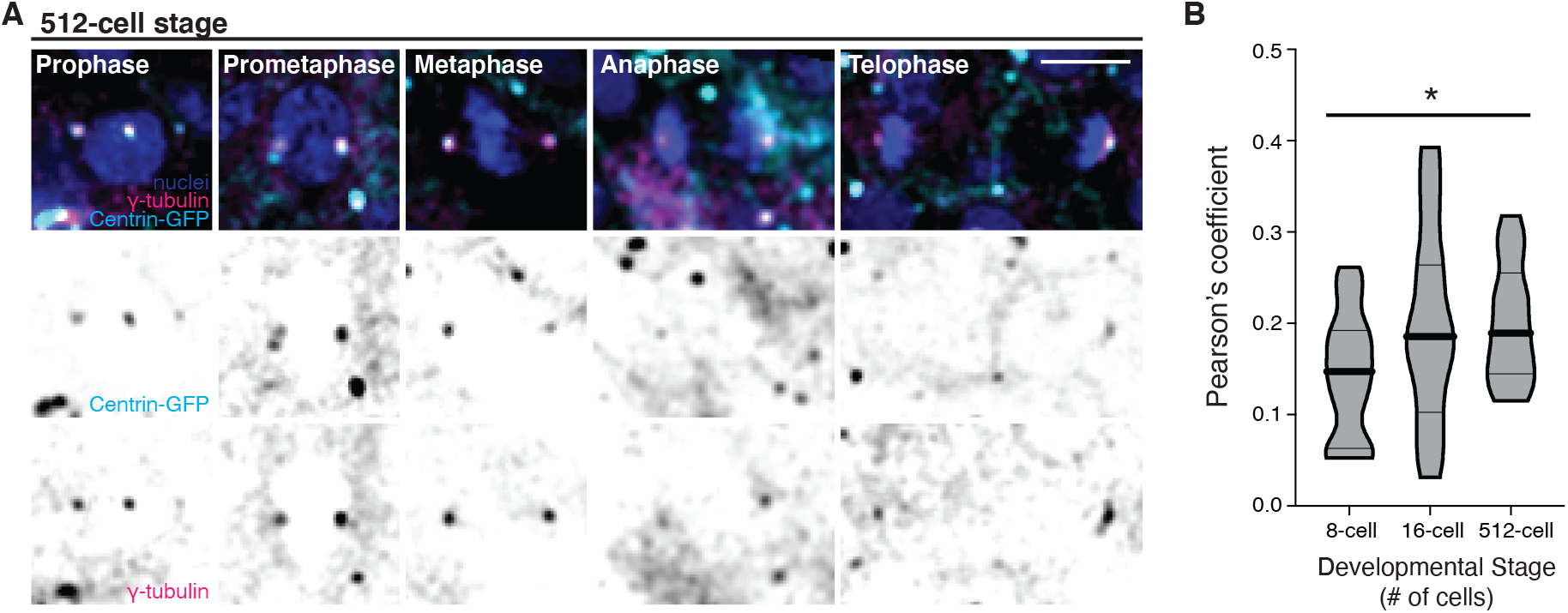
Centrosomes in early zebrafish development are uniquely structured. **(A)** Maximum confocal projections of fixed mitotic zebrafish cells at the 512-cell stage of development. Mitotic stages denoted. Centrin-GFP (magenta, inverted grayscale), γ-tubulin (cyan, inverted grayscale), and nuclei (DAPI, blue) shown. Bar, 10μm. **(B)** Violin plot depicting Pearson’s correlation coefficient between centrin-GFP and γ-tubulin signal in fixed zebrafish embryo cells at the 8- (n=23 embryos), 16- (n=19 embryos), and 512-cell stage (n=14 embryos). One-way ANOVA, p=0.0488 (*). Plot boundaries depict minimum and maximum, 25^th^ and 75^th^ quartiles represented by thin black line, median represented by thick black line. Detailed statistical analysis in Methods Tables.

**Supplementary Figure 3.**
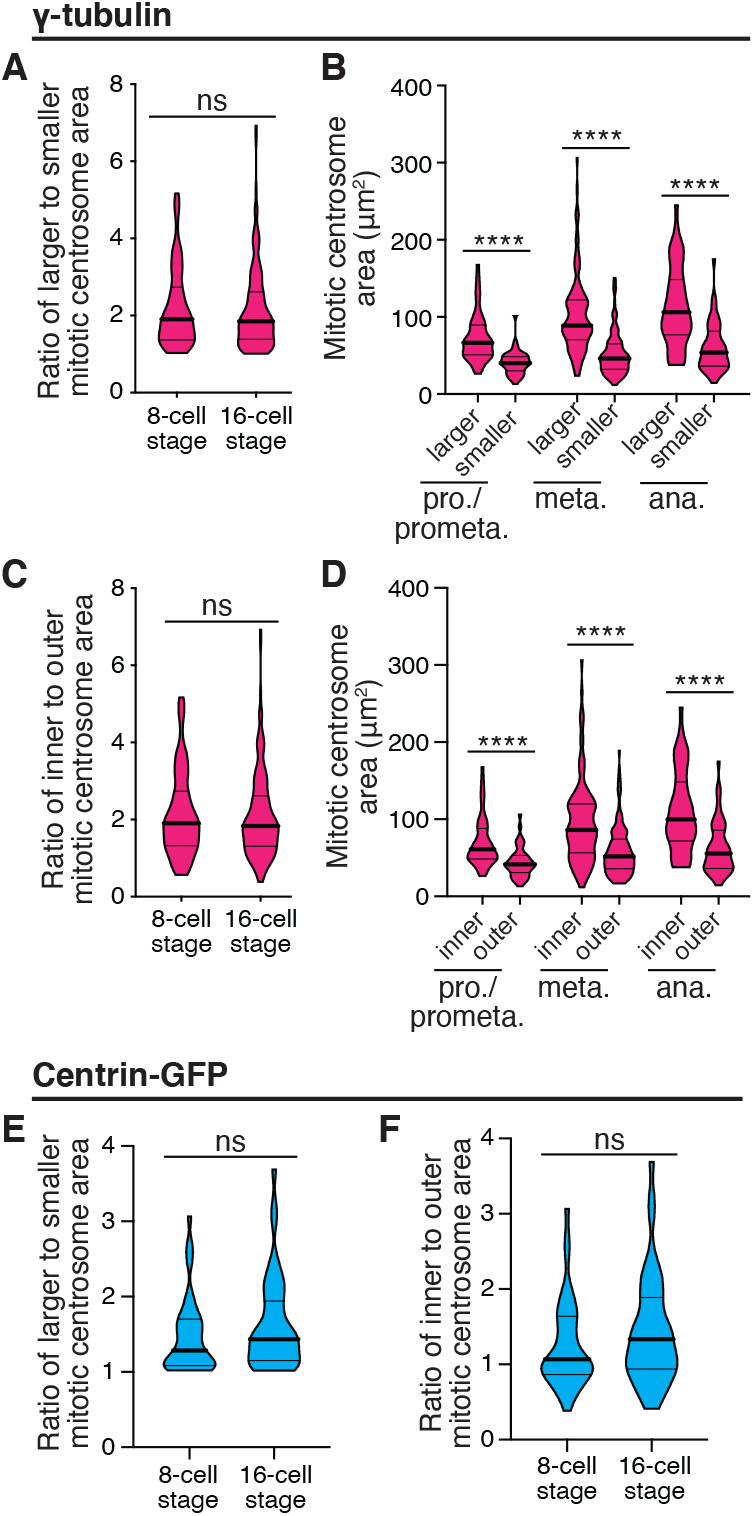
Mitotic centrosomes are asymmetric during early zebrafish cell divisions. **For graphs A-D:** measurements from γ-tubulin signal. **(A)** Violin plot depicting the ratio of mitotic centrosome areas binned by size (larger-to-smaller centrosome ratio. Student’s t-test, p=0.7101 (ns). **(B)** Mitotic centrosome area of 16-cell stage embryo during prophase/prometaphase, metaphase, and anaphase binned by size (larger or smaller). Student’s t-test, p<0.0001 (****). **(C)** Violin plot depicting the ratio of mitotic centrosome areas binned by position in relation to the midline (inner-to-outer centrosome ratio). Student’s t-test, p=0.7221 (ns). **(D)** Mitotic centrosome area of 16-cell stage embryo during prophase/prometaphase, metaphase, and anaphase binned by position in relation to midline (inner or outer). Student’s t-test, p<0.0001 (****). **For graphs E-F:** measurements from centrin signal. **(E)** Violin plot depicting the ratio of mitotic centrosome areas binned by size (larger-to-smaller centrosome ratio. Student’s t-test, p=0.2015 (ns). **(F)** Violin plot depicting the ratio of mitotic centrosome areas binned by position in relation to the midline (inner-to-outer centrosome ratio). Student’s t-test, p=0.2063 (ns). **For all violin plots:** Plot boundaries depict minimum and maximum, 25^th^ and 75^th^ quartiles represented by thin black line, median represented by thick black line. Detailed statistical analysis in Methods Tables.

**Supplementary Figure 4.**
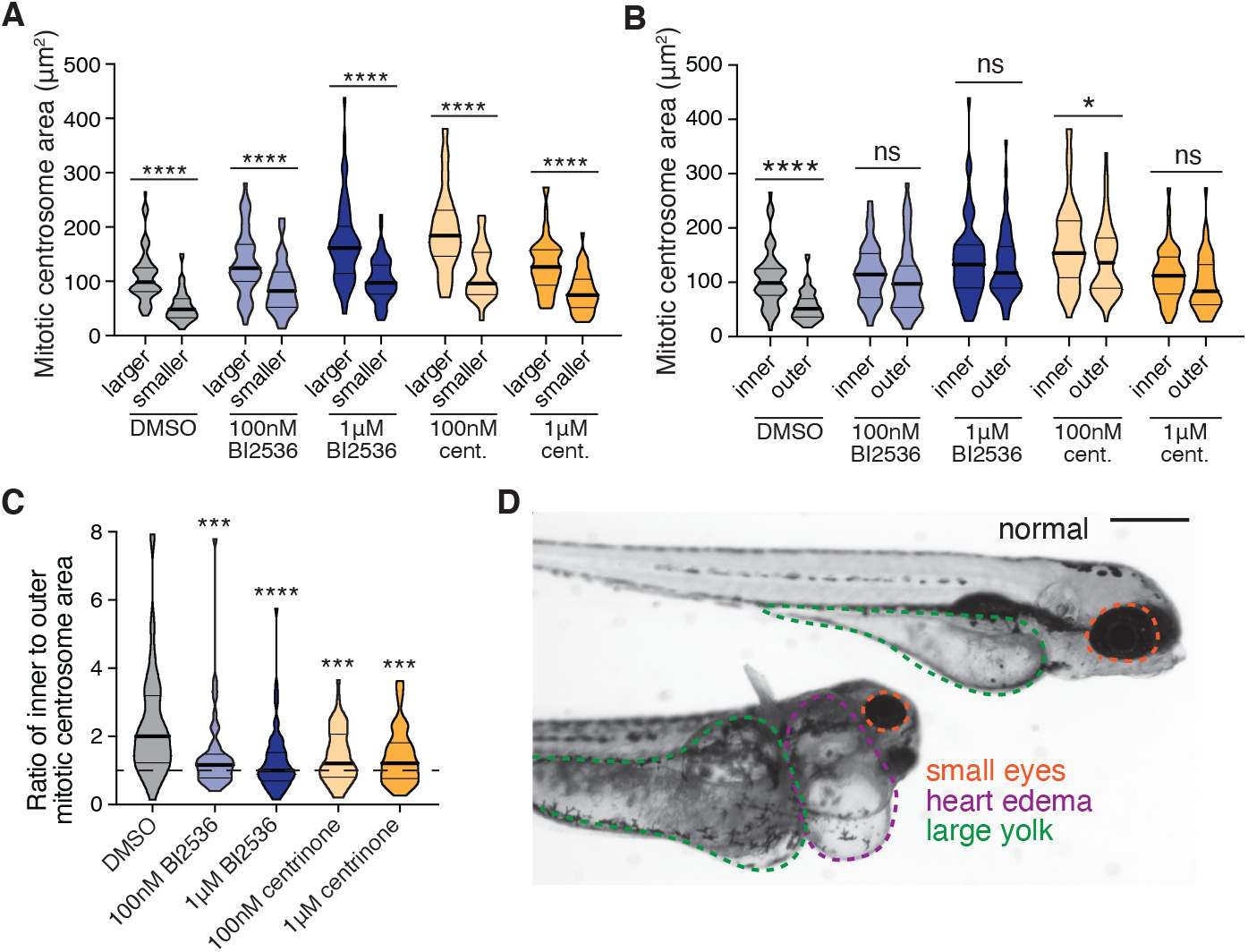
PLK1 and PLK4 activity are required for asymmetric mitotic centrosome positioning. **(A-B)** Violin plot depicting mitotic centrosome area under conditions of DMSO (gray), BI2536 (100nM or 1μM, blue), or centrinone (100nM or 1μM, gold) exposure. Mitotic centrosome area measured with γ-tubulin antibody signal and binned by size (larger/smaller in **A**) or position relative to midline (inner/outer, **B**). Student’s t-test performed within each treatment group. **(C)** Violin plot depicting the ratio of mitotic centrosome areas binned by position (inner-to-outer centrosome ratio) under conditions of DMSO (gray), BI2536 (100nM or 1μM, blue), or centrinone (100nM or 1μM, gold) exposure. Mitotic centrosome areas measured from γ-tubulin signal from fixed zebrafish embryos at the 16-cell stage. One-way ANOVA with Dunnett’s multiple comparison test. **(D)** Representative image of normal zebrafish (top) and abnormal zebrafish (bottom) displaying heart edema (purple), large yolk (green), and small eyes (orange) phenotypes that were quantified in Figure 4F. **For all graphs:** Violin plot boundaries depict minimum and maximum, 25^th^ and 75^th^ quartiles represented by thin black line, median represented by thick black line. Detailed statistical analysis in Methods Tables.

## Movie Legends

**Movie 1: Zebrafish and *C. elegans* embryogenesis.** Timelapse imaging of a zebrafish embryo (top) and *C. elegans* embryo (bottom) during the first several rounds of cell division. Microtubules (cyan), cell membrane (magenta), and nuclei (magenta) depicted. Bars, 50μm.

**Movie 2: Spindles position parallel to yolk boundary in zebrafish embryo divisions.** Timelapse imaging of EMTB-3xGFP transgenic zebrafish embryo shown with depth-coding. Cellular monolayer visualized from top of the embryo, yolk is behind cell layer in movie. Movie acquired over approximately 45 minutes.

**Movie 3: Live imaging of microtubules and centrosomes in zebrafish embryos.** Timelapse movie of EMTB-3xGFP (left) and centrin-GFP (right) transgenic embryos from the 8-cell to 32-cell stage of development. Images acquired over approximately 45 minutes for both movies.

**Movie 4: Single spindle dividing with microtubule and centrosome markers.** Single spindle depicted during a cell division in EMTB-3xGFP (inverted grayscale, left) or centrin-GFP (fire LUT, right). Images acquired over approximately 8 minutes (EMTB-3xGFP) and 12 minutes (centrin-GFP), respectively.

